# Expanding the family of genetically encoded voltage indicators: Insights from heliorhodopsins

**DOI:** 10.1101/2023.03.27.534354

**Authors:** Srividya Ganapathy, Xin Meng, Delizzia Mossel, Mels Jagt, Daan Brinks

**Affiliations:** Department of Imaging Physics, Delft University of Technology, Delft, The Netherlands; Department of Pediatrics & Cellular and Molecular Medicine, UCSD School of Medicine, San Diego, USA; Department of Molecular Genetics, Erasmus University Medical Center, Rotterdam, The Netherlands

**Keywords:** Voltage sensors, neuroscience, fluorescence microscopy, rhodopsins, protein engineering

## Abstract

Genetically encoded voltage indicators (GEVIs), particularly those based on microbial rhodopsins, are gaining traction in neuroscience as fluorescent sensors for imaging voltage dynamics with high-spatiotemporal precision. Here we establish a novel GEVI candidate based on the recently discovered subfamily of the microbial rhodopsin clade, termed heliorhodopsins. We discovered that upon excitation at 530-560nm, wild type heliorhodopsin exhibits near infra-red fluorescence which is sensitive to membrane voltage. We characterized the fluorescence brightness, photostability, voltage sensitivity and kinetics of wild type heliorhodopsin in HEK293T cells and further examined the impact of mutating key residues near the retinal chromophore. The S237A mutation significantly improved the fluorescence response of heliorhodopsin by 76% providing a highly promising starting point for further protein evolution.

## 1. Introduction

Detailed studies of neural circuitry and computation are contingent upon resolving the electrical dynamics of several neurons in parallel with high spatio-temporal precision. Direct visualization of changes in neural membrane potential has been facilitated by engineering bright and sensitive probes of which the fluorescence is modulated by changes in membrane voltage. These engineered transmembrane proteins are termed genetically encoded voltage indicators (GEVIs)^1^. Various GEVI families have been optimized over the past years and particularly GEVIs based on microbial rhodopsin proton-pumps have enabled the recording of activity in an ensemble of neurons with sub-millisecond response time^2^.

The first rhodopsin-based GEVI was derived from the bacterial Proteorhodopsin, discovered due to the success of metagenomic sequencing efforts in Monterey Bay^3^. Another proton-pump, Archaerhodopsin-3 (Arch) from the archaea Halorubrum sodomense, was found to be a better GEVI candidate for expression in mammalian cells^4^. The first Arch versions were very dim and required several iterations of molecular evolution^4–8^. Many flavours of Arch-based GEVIs have since been developed with improved brightness, sensitivity and membrane targeting, the most recent ones being Archon1 and Quasar6^4,9,10^. Quasar6a has a reported voltage sensitivity of 73±8% per 100 mV in HEK293T cells and a significant improvement in SNR in neurons over earlier versions^10^. The evolved brightness of Archon1 and Quasar6a have enabled in vivo imaging in mice and zebrafish, in combination with a spectrally orthogonal Channelrhodopsin for *in vivo* all-optical electrophysiology^9–11^.

Arch-based GEVIs exhibit complex photophysics and various models have been proposed over time to shed light on its voltage sensitivity^12,13^. Wild type Arch and some other rhodopsins typically display weak fluorescence arising from the retinal chromophore^14^. Retinal is covalently bound to the protein via a Schiff-baselinkage with a Lysine, which is normally protonated. The near infra-red fluorescence of this retinal protonated Schiff-base (RPSB) is modulated by the charge distribution of nearby residues lining the binding pocket. Light absorption initiates the photocycle of the protein via a sequence of conformational changes which in turn can impact RPSB fluorescence due to changes in electrostatic interactions. Canonically, photon absorption in the ground state leads to isomerization of the RPSB from *all*-trans to 13-cis and relocation of its proton to a negatively charged counterion acceptor (M-state). Photophysical characterization of Arch suggests that the reprotonation of the Schiff-base (M→N) is influenced by membrane voltage and populates the N-state, where an increased likelihood of photon-absorption leads to a fluorescent Q-intermediate^12^.

The complex photophysics of Arch and the high tunability of its fluorescent brightness, voltage sensitivity and kinetics by targeted mutations have made it an exciting candidate to investigate and evolve further as a GEVI^4,7,9,15,16^. However, besides Arch only a handful of rhodopsin proton-pumps have been engineered as GEVIs, despite the expansive diversity of the microbial rhodopsin family. Other rhodopsins with different ionic transport or sensory functions remain vastly unexplored as potential GEVIs, despite all having the same tunable retinal choromophore in common. In addition, novel rhodopsins with unique properties are continuously being added to the family, which deserve further exploration of their bioengineering potential. Recently, metagenomic sequencing in Lake Kinneret led to the discovery of a new family of rhodopsins termed Heliorhodopsins^17^. They were found to be abundant in the photic zone occurring in diverse host species ranging from bacteria to viruses^17^.

Heliorhodopsins are also heptahelical retinal binding proteins, but are remarkably different from other microbial rhodopsins due to an inverted insertion in the membrane with a cytoplasmic N-terminal^17^. Their precise physiological functions are thought to be diverse and are unknown as of yet. No clear ion translocation has been found, with the exception of a viral heliorhodopsin which functions as a light-gated proton channel^18^. They display a relatively long photocycle (∼1-5 seconds)^17,19^ indicating that they may have some kind of sensory or signaling role. Recent studies suggest that heliorhodopsins may be involved in membrane signaling via light-induced lipid remodelling^20^ or the transport of membrane-impermeable molecules^21^. The crystal structures of two heliorhodopsin variants have recently been resolved, shedding some light on their unusal properties^19,22^. Bacterial HeR-48C12 contains a large cytoplasmic RPSB cavity with several polar residues and water molecules. This arrangement enables transient proton-transfer from the RPSB and back, via a proton accepting group (involving H23 and H80). This polar H-bonded environment is highly amenable for tuning the spectral properties of retinal^23^, making Helios an interesting candidate for bioengineering.

In this study we demonstrate the potential of Heliorhodopsin (bacterial HeR-48C12, Figure 1a) to function as a fluorescent indicator of membrane voltage. We show that wild-type Heliorhodopsin displays voltage-dependent fluorescence, which can be improved with targeted mutations in the retinal binding pocket. This research paves the way for further evolution of Heliorhodopsin-based GEVIs and opens the door for engineering other members of the microbial rhodopsin clade.

**Fig. 1.**
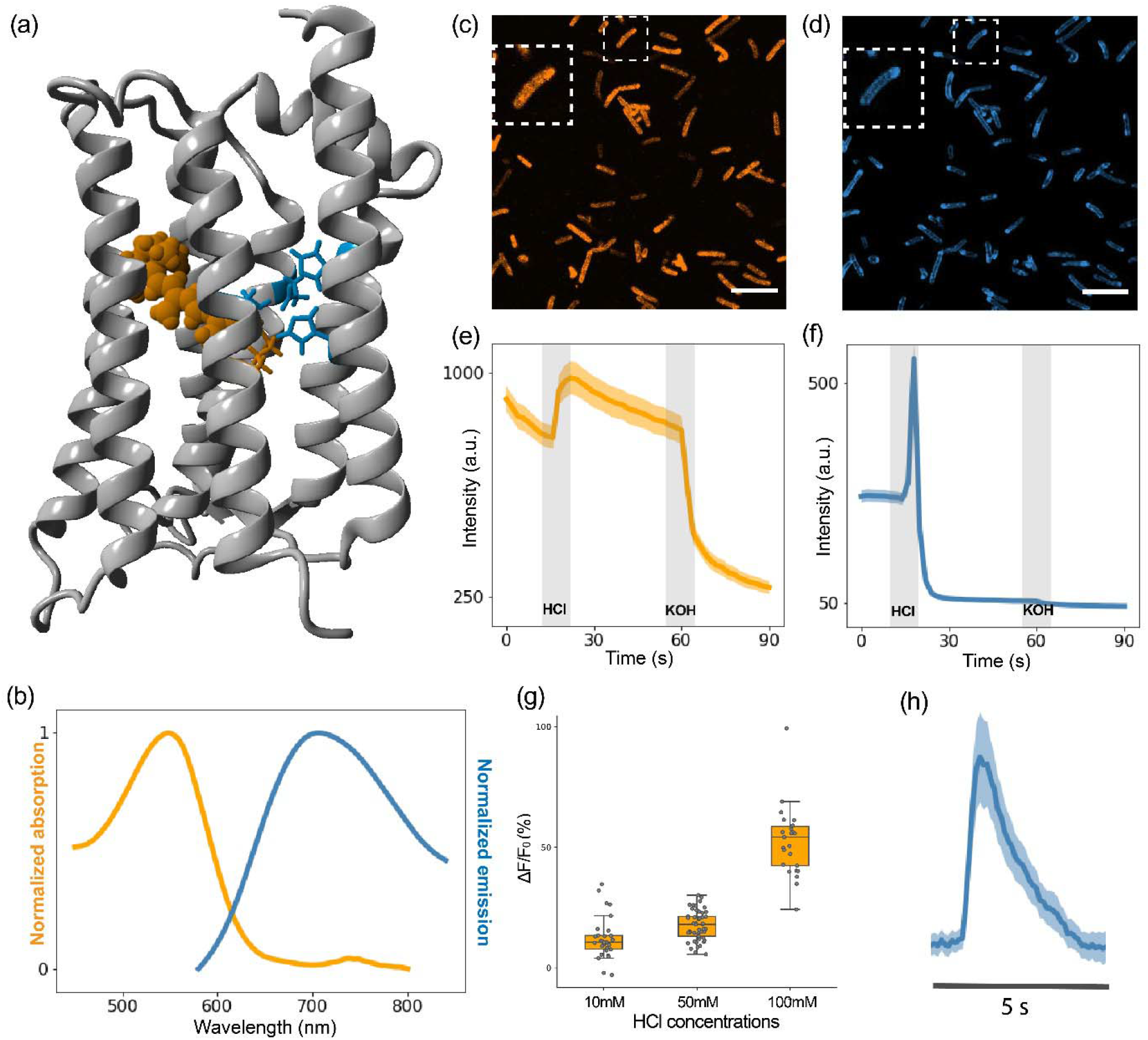
Preliminary characterization of Helios fluorescence. (a) Crystal structure of Helios48C12 (PDB 6su3) displaying the retinal Schiff-base in orange and residues involved in colour tuning in blue (b) Normalized absorption and emission spectra of purified WT Helios (c-d) Representative confocal fluorescence images of E. coli expressing Helios under 561 nm (c) and 640 nm (d) illumination, with the inset representing a zoom-in of an individual cell (e-f) Fluorescence response of E. coli expressing Helios to 25 mM HCl and 25 mM KOH addition under 561 nm (e) and 640 nm (f) illumination (n=45 cells). Videos were recorded at 1fps. The thick line is the mean response with the lighter region representing the SEM, the grey blocks indicate the time point of HCl or KOH addition. (g) Quantification of E. coli fluorescence response under 561 nm illumination to increasing concentrations of extracellular HCl addition (10 mM: mean value = 12.23; n=35; 50 mM: mean value = 17.55; n=50; 100 mM: mean value = 52.28; n=39; in the boxplots the boundaries of the whiskers are based on an interquartile range of 1.5, each grey dot in the boxplot represents a cell. (h) Spinning-disk confocal fluorescence response of Helios at 640 nm illumination recorded at 10 fps. Mean and SEM as in (e-f)

## 2. Materials and Methods

### 2.1. Plasmids and materials

The pBAD vector for recombinant expression of Heliorhodopsin HeR-48C12 containing an N-terminal 6xHis tag (pBAD-Helios-NT-6xHis) was a kind gift from Alina Pushkarev and Oded Béjà. The sequences for targeting and endoplasmic reticulum export motifs (TSX3ER2) and Citrine were derived from MPC020: CamKII CMV_NovArch_citrine, which was a gift from Adam Cohen (Addgene plasmid # 153193)^8^. The pCAG backbone was derived from pCAG-Archon1-KGC-EGFP-ER2-WPRE, which was a gift from Edward Boyden (Addgene plasmid # 108423)^9^.

### 2.2. *E.coli* culturing and purification

pBAD-Helios-NT-6xHis was transformed into chemically competent *E. coli* cells (NEB® 5-alpha, NEB) as per the manufacturer’s protocol. Overnight cultures were grown in LB medium under ampicillin selection (100 µg/mL) in a shaking incubator at 37°C, 150 rpm. The following day, the culture was diluted 1:50 times to a volume of 400 mL. Opsin expression was induced an OD_600_ of 0.4-0.6 by adding a final concentration of 0.2% arabinose. 20 µM all-trans retinal dissolved in ethanol was added to the culture and it was left shaking for another 14-18 hours. The cells were pelleted by centrifugation at RT, 4000g, 20 min and washed twice with an equal culture volume of 150 mM NaCl. The cell pellet was resuspended in 4 mL lysis buffer containing 50 mM Tris, 300 mM NaCl, 0.1% DDM and lysed using a French press. Membrane vesicles were pelleted by ultracentrifugation at 100,000g, 45 min, 4°C. The pellet was resuspended in 50 mM Tris, 20 mM imidazole, 300 mM NaCl, 2% DDM, pH 6.5 and was left to mix for 1 hour, RT. Insoluble debris was spun down at 100,000g, 45 min, 4°C. and the supernatant was loaded onto a column containing Ni^2+^NTA resin for purification of the His-tagged protein using affinity chromatography. The resin was washed with 10 bed volumes of 50 mM Tris, 50 mM imidazole, 300 mM NaCl, 0.1% DDM, pH 6. The purified protein was eluted in 50 mM Tris, 500 mM imidazole, 300 mM NaCl, 0.1% DDM, pH 6 and concentrated using a 10 kDa spin column (Millipore). The absorption and emission spectra of the purified protein were recorded at RT (Lambda365, Perkin Elmer and FLS980, Edinburgh Instruments)

### 2.3. Confocal imaging

*E. coli* cells expressing pBAD-Helios-NT-6xHis were grown as described above. The cell were spun down and the cell pellet was washed thrice and resuspended in an equal culture volume of PBS. The cell suspension was plated onto 35 mm imaging dishes with a 10 mm glass coverslip (Cellvis) coated with poly-L-lysine (Thermo-Fisher). Fluorescence images were captured using laser-scanning confocal microscopy (Nikon Eclipse Ti inverted) at excitation wavelengths of 561 and 640 nm and emission at 595/50 nm and spinning disk (IX81, Olympus) confocal microscopy

### 2.4. Cloning

HeR-48C12 was amplified from pBAD-Helios-NT-6xHis and combined with TSX3ER2 and Citrine using overlap-extension PCR with Phusion high fidelity master mix (NEB). The primers used for cloning are listed in the Supporting Information (SI, Table S1). TX3ER2 and Citrine were inserted at the C-terminal end of the protein, due to the inverted orientation of Heliorhodopsin in the membrane. The pCAG backbone was amplified using a high fidelity polymerase KODextreme hot start (Merck Sigma). TSX3ER2-Citrine-Helios was inserted into pCAG using Gibson assembly (Gibson assembly mastermix, NEB) to generate pCAG-Helios. Point mutations were generated by PCR using end-to-end primers with the mutation site encoded in the forward primer. KODextreme polymerase was used for amplification and the product was ligated (KLD enzyme mix, NEB) and transformed into NEB® 5-alpha Competent cells. The primer sequences used for mutagenesis are provided in the supplementary information (SI, Table S1).

### 2.5. HEK cell culturing

HEK293T cells were grown at 37°C, 5-10% CO_2_ in Dulbecco’s modified Eagle medium supplemented with 10% fetal bovine serum and Penicillin/Streptomycin. The cells were transfected at a confluency of ∼80% with 600-1000 ng of plasmid and 6 uL of TransIT293T (Mirus) transfection reagent. 24 hours after transfection, the cells were plated onto 35mm imaging dishes containing a 10 mm glass coverslip (Cellvis) coated with fibronectin (Merck).

### 2.6. Patch clamp electrophysiology

Whole cell voltage clamp recordings were performed at room temperature (25°C) 48-72 hours after transfection. The cells were rinsed with extracellular buffer containing 125 mM NaCl, 2.5 mM KCl, 15 mM HEPES, 1 mM CaCl_2_, 1 mM MgCl_2_ and 30 mM Glucose; pH 7.3; osmolarity adjusted to 310 mOsm. Micropipettes were pulled from borosilicate glass capillaries (World Precision Instruments, 1.5 mm OD, 0.84 mm ID) using Next Generation Micropipette Puller (Sutter Instrument, P-1000) to obtain a pipette resistance of 5-10 MΩ. The pipettes were filled with intracellular buffer containing 125 mM potassium gluconate, 8 mM NaCl, 0.1 mM CaCl_2_, 0.6 mM MgCl_2_, 1 mM EGTA, 10 mM HEPES, 4 mM Mg-ATP and 0.4 Na-GTP; pH 7.3; osmolarity adjusted to 295 mOsm. The micropipettes were positioned using the Patchstar micromanipulator (Scientifica). All patch clamp data were acquired in voltage-clamp mode using AM Systems Model 2400 Patch Clamp Amplifier, using voltage steps ranging from -100 to 100 mV. Simultaneous fluorescence measurements were performed as described below.

### 2.7. Fluorescence imaging

These experiments were performed using a home-built multi modal microscope with a patch-clamp add on, the design of which has been described recently^24^. Epi-fluorescence imaging was performed using two laser beams at 488 nm (OBIS 488 LX, Coherent) and 532nm (MLL-III-532, CNI) focussed onto the back aperture of a 25X objective (XLPLN25XWMP2, Olympus). An illumination power of 11.2 mW/mm^2^ and 37.23 mW/mm^2^ were used for brightness screening for 488nm and 532 nm respectively. For patch clamp characterization, the 532 nm intensity was 87.6 mW/mm^2^. The emission light was filtered using a multi-band dichroic mirror (Di03-R405/488/532/635-t3-32×44, Semrock) and a 552-779.5 bandpass filter (FF01-731/137-25, SemRock). The images were acquired at a frame rate of 100 or 500 Hz using an sCMOS camera (ORCA Flash4.0 V3, Hamamatsu; 2048 × 2048 pixels, 6.5 µm pixel size). The voltage pulses, illumination and camera recording were synchronized using the National Instruments DAQ (USB-6363). All software for controlling the hardware, image acquisition and analysis were custom written in Python^24^.

### 2.8. Data analysis

From the recorded camera video, a background region was manually selected. The averaged background trace from this region was calculated, then smoothed. At each frame, time-corresponding background value was subtracted. A maximum-likelihood pixel weighting algorithm was utilized to extract the fluorescence trace. The trace was then corrected for photo-bleaching by normalization against the bi-exponential fitting of itself. An averaged period was calculated from it. To measure the change of fluorescence in response to membrane voltage, *F*_*bl*_ (baseline fluorescence) and *F*_*ss*_ (steady-state fluorescence) values were computed from fluorescence at resting potential and during voltage step after reaching steady state, then the sensitivity ΔF/F was presented as (*F*_*ss*_ - *F*_*bl*_)/*F*_*bl*_. To show the protein dynamics, the up-swing and down-swing phases were segmented, then fitted to a bi-exponential function: 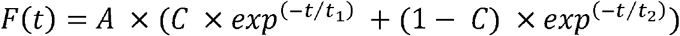, in which A is a constant, C is the magnitude percentage between two single exponential functions, *t*_l_ is the fast time constant and *t*_2_ is the slow time constant.

## 3. Results and Discussion

### 3.1. Expression and fluorescence imaging of Helios in *E. coli*

In order to assess the fluorescence properties of wild-type Heliorhodopsin (hereby called Helios), we did a preliminary characterization in *Escherichia coli*. Recombinant N-terminal 6XHis tagged Helios was overexpressed in E. coli and purified using Ni^2+^ NTA affinity chromatography. The absorption spectrum of purified Helios displayed a λ_max_ at 549±1 nm, in good agreement with reported values^17^. Upon excitation at 550nm, we obtained a distinct emission band extending from 600 nm into the near infra-red region, peaking at ∼700nm (Figure 1b). Based on comparison of absorbance and emission integral to fluorophores with known quantum yield, we estimate a fluorescence quantum yield of 6×10^−4^ which is in agreement with the typical range reported for other microbial rhodopsins^4,5,25^.

Next, we directly imaged Helios in intact E. coli cells using confocal microscopy. Bright fluorescence was seen localized to the cell membrane when imaged under 561 and 640 nm illumination (Figure 1c and 1d), though with some moderate photobleaching (SI, Figure S1). Biexponential fitting of the photobleaching response yielded time constants of 20 and 120 s for the 561 and 640 nm fluorescence respectively (SI, Figure S1). We tested the sensitivity of this fluorescence to changes in extracellular pH, as a preliminary indicator of voltage sensitivity. Upon addition of 25 mM HCl to the cells, an increase in fluorescence was seen in the 561 nm channel well above the photobleaching background (Figure 1e). This step response in fluorescence was roughly linear with increasing concentrations of HCl (Figure 1f) and could be reversed upon addition of 25 mM KOH (Figure 1e). However, in the 640 nm channel, we observed an initial large rise in fluorescence followed by rapid quenching of the signal within 3-4 seconds (Figure 1g,h). This fluorescence could not be recovered with dark incubation or KOH addition (Figure 1g). Furthermore, it was not impacted by the presence or order of the 561 nm illumination pulse. This quenching reaction possibly involves a complex photocycle pathway characterized by the pH dependent inactivation of a near infra-red photointermediate. Since our interest is in the use of Heliorhodopsin as GEVI which requires a linear and reversible response to membrane voltage, we focused on the ∼561 nm fluorescence of Helios.

### 3.2. Characterization of Helios WT in HEK293T

Helios was cloned into an expression vector for human embryonic kidney (HEK293T) cells driven by the strong pCAG promoter. Based on prior efforts to optimize the membrane trafficking of Arch^4^, we added a tri-repeat of targeting sequences and endoplasmic reticulum motif (TSX3ER2) with Citrine as a fusion protein for localization (Figure 2a), based on the design of Quasar3^26^. HEK293T expressing pCAG-Helios was imaged using a home-built epi-fluorescence microscope with a patch-clamp add on for electrophysiology (Figure 2b). Strong near infra-red fluorescence (660–800 nm) could be seen when imaged at 488 and 532 nm in agreement with the measurements in *E. coli* (Figure 2c). Interestingly, no fluorescence was seen upon 639 nm excitation, possibly due to differences in binding-pocket conformation in the mammalian expression system versus E. coli. While Helios expression was mostly localized to the plasma membrane (Figure 2c), intracellular aggregates and overexpression leading to cell death were also seen, likely due to the strong pCAG promoter. The ratios of membrane fluorescence to soma fluorescence, as measured by quantifying the image intensity at the cell contour and soma respectively, show similar values around 1 for WT and mutants, indicating that expression is distributed within the cell (SI, Figure S3). Helios exhibited moderate photobleaching at 532 nm (SI, Figure S2), with a fast constant of 11.16 ± 3.39 ms (54.9% ± 0.14%) and a slow constant of 158.39 ± 3.39 ms (n = 6 cells; all statistics are mean ± s.t.d.).

**Fig. 2.**
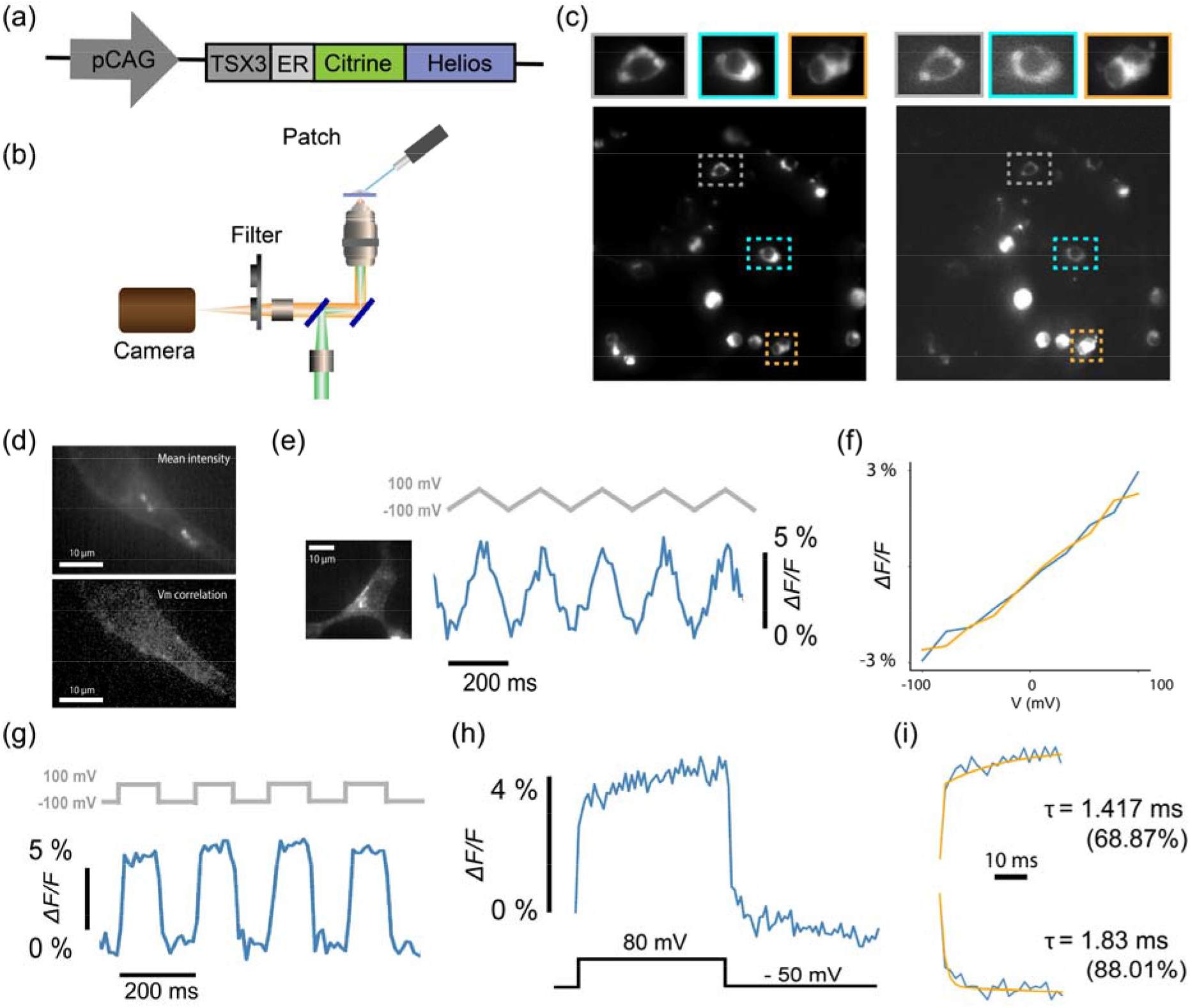
Characterization of voltage sensitivity of WT Helios in HEK293T cells. (a) Schematic of the plasmid for expression of Helios under the pCAG promoter with the targeting motifs (TSX3, ER) and Citrine as a fusion protein (b) An illustration of the setup used for simultaneous fluorescence imaging with voltage-clamp electrophysiology (c) Full field of view fluorescence images of HEK293T cells expressing Helios WT under 488 nm (left) and 532 nm excitation (right). The zoom-in view on the top displays representative individual cells. (d) Top: mean intensity image from a video of a voltage-clamped cell expressing Helios. Bottom: The correlation map between the video and the membrane voltage. (e) Characterization of the fluorescence response of Helios to whole cell voltage-clamp. Left: Fluorescence image of the patched HEK293T cell. Right: Helios fluorescence response to 200 mV voltage ramps recorded at 100 fps. The illumination intensity was 87.6 mW/mm^2^. (f) Averaged upswing and downswing traces from 25 trials. (g) Helios fluorescence response to 200 mV voltage steps recorded at 100 fps. (h) Averaged fluorescence response to 130 mV voltage steps recorded at 500 fps. (e) Biexponential fitting analysis and kinetics of voltage sensitive fluorescence.

We assessed the voltage sensitivity of Helios at room temperature by modulating the membrane potential of HEK293T cells expressing Helios using whole cell patch-clamp electrophysiology and measuring the changes in fluorescence. Correlation of per-pixel fluorescence change with the change in membrane voltage showed characteristic localization of the voltage sensitive fluorescence at the cell membrane (Figure 2d). In combination with mean intensity images displaying significant somatic fluorescence (Figures 2c,d), this indicates Helios displayes proper membrane trafficking but is overexpressed in most cells. 200mV voltage ramps where used at a frequency of 5 Hz (Figure 2e). The concurrent fluorescence response was recorded at 532 nm on a sCMOS camera at a frame rate of 100 Hz. Helios displayed a linear response to membrane voltage over a -100 to +100 mV range (Figure 2f). Subsequently, 200 mV voltage pulses were delivered at a frequency of 5 Hz, for a total duration of 5 seconds. Here, the 532 nm fluorescence response was recorded at a frame rate of 100 Hz (Figure 2g) or 500-1000 Hz for high-speed characterization of the time constants (Figure 2h,i). The signal was temporally averaged after subtracting the background signal and correcting for photobleaching. The fractional change in fluorescence was extracted and normalized to a 200 mV step (from -100 mV to +100 mV) yielding a ΔF/F_0_ of 6.14% ± 1.35% per 200 mV for Helios WT (n = 7 cells; mean ± s.t.d.) Biexponential fitting of the fluorescence trace measured at 500 Hz yielded a fast time constant for the upswing of 2.06± 0.47 ms (62%) and of 2.40±0.40 for the downswing (n = 3 cells). (Figure 2i).

### 3.3. Comparison between Helios mutants

The voltage response of Helios appeared to be substantially dominated by the recording speed in our measurements. We were intrigued by the step-like fluorescence response to the 200 mV voltage block recorded at 500 Hz, since this is a compable speed at which in vivo voltage imaging is typically performed where Arch-based sensors tend to have comparable or slower responses. Therefore, we attempted to improve the fluorescence and voltage sensivity of Helios by targeted mutations in the retinal binding pocket.

Arch and other rhodopsin proton-pumps typically have two negatively charged counterion resides functioning as a complex. In contrast, Helios contains a single E107 as the counterion, which is hydrogen bonded to the RPSB along with an uncharged S237 (Figure 3a)^17,22^. E107 has a pKa at 3.7 and is therefore likely to be unprotonated under physiological conditions^4^. The counterion influences the charge distribution of the RPSB, and mutations to neutral residues often cause spectral red-shifts in many rhodopsins^27^. In Arch, the red-shifting D95N and D95Q mutations eliminated the photocurrent and improved the voltage sensitivity^7,15^. Thus, we tested the analogous E107N and E107Q mutations in Helios as a first target. However, the E107N and E107Q mutants showed diminished fluorescence (Figure 3b,c) with the WT brightness being 3 times higher than E107N and 5 times higher than E107Q (23 E107N and 107 E107Q cells contribute to the mean brightness calculation; 532 nm, 87.6 mW/mm^2^). No clear voltage response could be measured for these mutants.

**Fig. 3.**
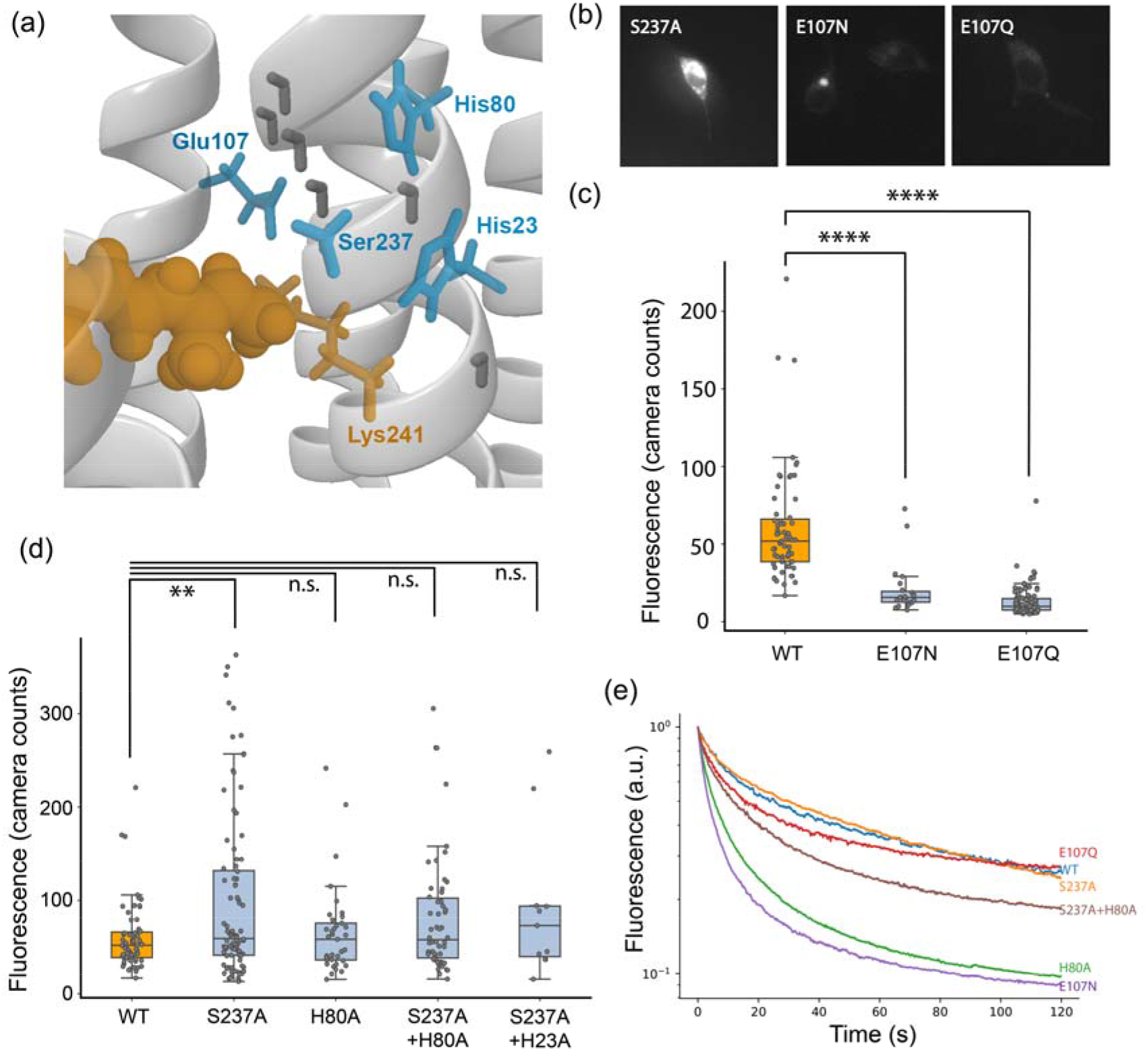
Brightness and photobleaching of Helios mutants. (a) Binding pocket view of the crystal structure of Helios displaying the retinal Schiff base (orange) and key colour tuning residues (blue). (b) Representative fluorescence images of HEK293T cells expressing the Helios mutants S237A, E107N and E107Q. The contrast across the images is adjusted to the same level. (c) Comparison of brightness values of HEK293T cells expressing Helios WT (mean value = 60.6 ± 36.1, n = 63 cells), E107N (mean value = 20.4 ± 15.9, n = 23 cells) and E107Q (mean value = 12.7 ± 9.0, n = 107 cells). The p-values of independent t-tests against Helios WT are: E107N :1.7e-6; E107Q: 2.4e-27. In the boxplots in subfigure (c) and (d), the boundaries of the whiskers are based on an interquartile range of 1.5, and each grey points in the plot represents one measurement. All statistics are mean ± s.t.d. (d) Comparison of brightness values of HEK293T cells expressing WT Helios (mean value = 60.6, n = 63 cells), S237A (mean value = 97.8 ± 88.3, n = 92 cells), H80A mean value = 65.3 ± 47.4, n = 37 cells), S237A+H80A (mean value = 81.7 ± 63.9, n = 54 cells) and S237A+H23A (mean value = 91.4 ± 78.2, n = 11 cells). The p-values of independent t-tests against Helios WT are: S237A: 0.00186; H80A: 0.57; S237A+H80A: 0.027; S237A+H23A: 0.037. The illumination intensity was 37.23 mW/mm^2^. (e) Normalized photo bleaching traces of Helios WT and mutants.

This may be attributed to the stronger interaction of the Helios RPSB with the surrounding water dense Schiff base cavity^22^. In most microbial rhodopsins, the counterion usually functions as the primary acceptor for proton transfer from the RPSB upon isomerization^28^. However in heliorhodopsins, the Schiff base cavity collectively functions as the primary proton acceptor^17,19,22^. The hydrogen bonded network involving charged binding pocket residues and water molecules participates in transient transfer of the Schiff base proton from and back to the RPSB^17^. A recent study identified colour tuning mutations in several of these conserved binding pocket residues, including H23, H80 and S237, which interact directly with the RPSB^23^. We therefore turned our attention to these sites, focusing on the mutations which were reported to cause spectral red-shifts i.e. H23A, H80A and S237A^23^ (Figure 3a).

We screened HEK293T cells expressing the above mutants for their fluorescence brightness (532 nm, 37.23 mW/mm^2^). The averaged brightness of S237A is 23% higher than the WT (the mean fluorescence is calculated from 63 WT cells and 92 S237A cells, p-value=0.00186), while there were no significant change for the other tested mutations (Figure 3d). However, prolonged illumination of the cells revealed differences in photobleaching behavior among the Helios mutants. While S237A and E107Q had comparable photobleaching rates to the WT, H80A and E107N photobleached significantly faster. Bioexponential fitting of the photobleaching curves revealed that the photobleaching is of a different nature in WT compared to the mutants: where WT bleaching is characterized by a relatively strong, high and fast time constant, that of S237A is dominated by a relatively strong, but less high and slow time constant. (SI, Figure S2). In a combination mutant, S237A saved some of the long term fluorescence loss of H80A (Figure 3e).

### 3.4. Voltage sensitivity of Helios S237A mutants

Because of the increased fluorescence brightness of the S237A mutant and positive effect on photobleaching, we focused our investigations of voltage sensitivity on S237A and mutant combinations with it (Figure 4). We assessed the voltage sensitivity of S237A, S237A+H80A and S237A+H23A (Figure 4a). Sensitivities were (as ΔF/F per 200 mV): 6.14% ± 1.35% for Helios WT (n = 7 cells; all statistics are mean ± s.t.d.), 6.49% ± 1.4% for S237A (n = 4 cells), 5.11% ± 1.23% for S237A+H80A (n = 5 cells) and 5.86% ± 0.17% for S237A+H23A (n = 4 cells) (Figure 4b). No statistically significant difference in voltage sensitivity was measured between any of the mutants. We compared the response speed of the S237A mutant to a 200mV voltage step to that of Helios WT (Figure 4c). We found that S237A had a response time of 1.69± 0.04 ms (82%) (n = 2 cells) and 1.95± 0.11 (86%) for the up- and downswing respectively (Figure 4d). We found no significant difference in the speed of voltage response between S237A and WT (Figure 4e).

**Fig. 4.**
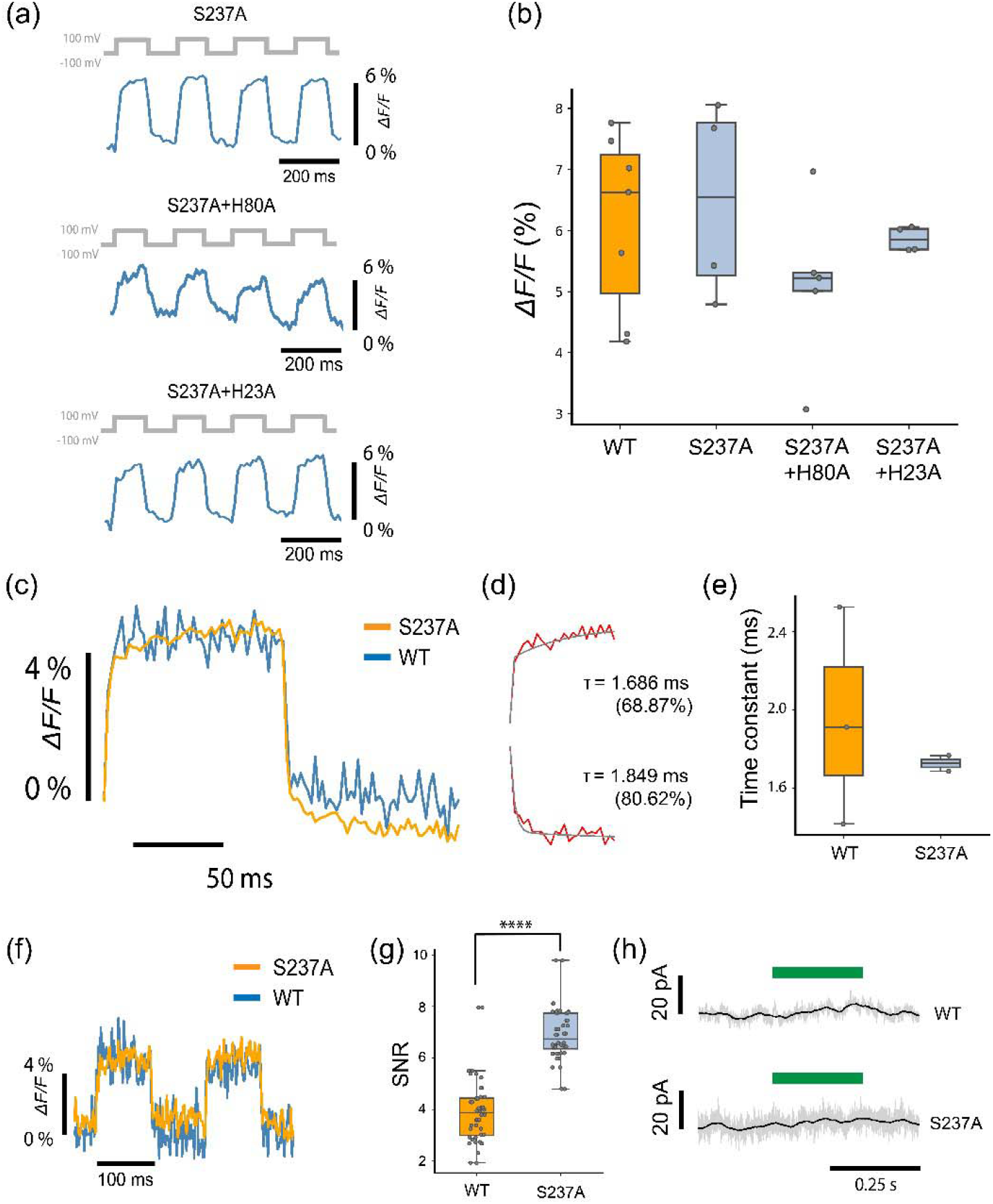
Characterization of voltage sensitivity of Helios mutants. (a) Fluorescence traces of Helios mutants in response to 200 mV voltage clamp square waves. From top to bottom: S237A, S237A+H80A and S237A+H23A. (b) Comparison of voltage sensitivity between Helios WT and mutants. The voltage sensitivities per 200 mV are: Helios WT: 6.14 ± 1.35 %; S237A: 6.48 ± 1.40 %; S237A + H80A: 5.11 ± 1.23 %; S237A + H23A: 5.86 ± 0.18 %. All statistics are mean ± s.t.d In the boxplots in subfigure (b), (e) and (g), the boundaries of the whiskers are based on an interquartile range of 1.5, and each grey point in the plot represents one measurement. (c) Overlay of averaged fluorescence response (25 periods) to 200 mV voltage steps from Helios WT and S237A, at 500 fps. S237A has fast kinetics as the upswing tau = 1.686 ms (fast component percentage 68.87%) and downswing tau = 1.849 ms (fast component percentage 80.62%) (d) Comparison between Helios WT and S237A rising fast time constants. Helios WT: 1.95± 0.45 ms (62%) (n = 3 cells); S237A: 1.72± 0.04 ms (82%) (n = 2 cells). All statistics are mean ± s.t.d. (e) Overlay of raw fluorescence response to a 200 mV voltage step from WT and S237A, at 500 fps. (f) SNR bar graphs of the WT (3.93 ± 1.3/500Hz, n = 48 measurements) and S237A (6.93 ± 1.0/500Hz, n = 48 measurements). The SNR from S237A is significantly higher (76.3%) than that from the WT. (g) Both the WT and S237A show no photocurrent upon 532 nm laser illumination. All the data shown here were acquired under an illumination intensity of 87.6 mW/mm^2^.

Given the relatively minor increase in brightness of S237A compared to WT and the similar voltage sensitivity and response speed, we were intrigued by the fact that the S237A fluorescence traces were substantially less noisy than the WT traces (Figure 4f). We quantified the signal to noise (SNR) with which we could detect a voltage step with Helios WT and S237A. We calculated the signal as the difference in the average fluorescence value for the 100 ms where the voltage was +100 mV, with the average fluorescence value for the 100 ms where the voltage was -100 mV. We calculated the noise as the standard deviation of the fluorescence for the 100 ms where the voltage was +100 mV. We found that the voltage detection SNR for S237A (6.93 ± 1.0/500Hz, n = 48 measurements) is 76.3% higher than that of WT Helios (3.93 ± 1.3/500Hz, n = 48 measurements). We wondered whether the increased noise in the WT recordings was due to photocurrent effects, but measured no discernible photocurrent at -30 mV upon illumination with green light (532 nm, 87.6 mW/mm2) in either WT or S237A (Figure 4h)

### 3.5. Comparison of voltage sensitive fluorescence between Helios and Arch

Voltage sensitive fluorescence is a property of the electrostatics of the retinal environment and accessibility to proton transfer via the hydrogen bonded network. The binding pocket and water cavity is quite different between Helios and Arch, which complicates a direct comparison. The Arch binding pocket contains 3 water molecules^30^, while for Helios at least 7 internal water molecules have been reported in the retinal cavity^31^. We will attempt to shed light on the discrepancy between the voltage sensitives of the Helios and Arch counterion mutants based on insights from the Helios crystal structure and mutation studies.

Prior work on Arch variants indicates that the increase in fluorescence quantum yield arises from the protonated Schiff base and a neutral counterion, where voltage regulates the equilibrium between protonated and deprotonated SB^12,32^. Mutation of the Arch counterion D95X destabilizes the protonated Schiff base leading to protonation and an increase in fluorescence under positive membrane voltage. However in Helios, E107 does not stabilize the protonated SB as effectively as in Arch. The weaker counterion interaction is compensated for by surrogate counterions involving other residues in combination with the water cavity and possibly anions. Unlike Arch, Helios has an S237 at the second counterion position (D222 in Arch). The large red shift of the mutant S237A indicates that this residue (in combination with surrounding water network) is probably crucial in stabilizing the charge on the protonated SB^23^.

Studies on the E107Q mutant indicate that reorganization of the retinal binding pocket stabilizes the protonated SB due to interactions with S237^33^ or even anions, as E107Q could bind anions even at physiological pH values^29,34^. Thus neutralizing the counterion in Helios (as in the E107Q mutant) is not analogous to the Arch D95X mutation. However our results with S237A demonstrate that this may be a useful fluorescence and/or sensitivity tuning site instead of E107, also due to evidence of its reorientation during proton transfer^22^.

Another important difference is that the primary proton transfer event in Helios occurs in the cytoplasmic direction, followed by proton back transfer to the RPSB. This could reduce the fidelity of deprotonation of the SB under negative voltage, thereby limiting the voltage sensitive response. A combination of mutations at S237 and/or other binding pocket residues which can stabilize the retinal protonated Schiff base will likely improve the voltage sensitive fluorescence of Helios.

## 4. Conclusion

We investigated the potential of Heliorhodopsin as a GEVI and the effect of several mutations on its brightness, voltage sensitivity, photobleaching statistics, response speed and photocurrent characteristics. The S237A mutant had a beneficial effect on fluorescence brightness without compromising photobleaching, voltage sensitivity or response speed and can be used as a template for further protein evolution. Since S237A is directly hydrogen bonded to the RPSB near H80 and is an important colour tuning residue, saturation mutagenesis of S237 or further mutant combinations in the binding pocket will likely yield improved variants. Additionally, membrane targeted expression of Helios variants can be improved by modifying the design of the expression construct, for instance by rearranging of trafficking motifs, using a different fusion protein or inserting spacer elements.

We expect that future electrophysiological investigations into Heliorhodopsins might increase our understanding of its native function and exact photodynamics, which will aid further bioengineering efforts.

## Acknowledgements

The authors would like to thank Alina Pushkarev and Oded Béjà from the Technion-Israel Institute of Technology for the pBAD heliorhodopsin plasmid used in this study. We gratefully acknowledge Leonie Gouweleeuw and Huma Safar for their advice and technical support provided during the course of this research, Marco Locarno for the help on computing the quantum yield. DB acknowledges support by an NWO Start-up Grant (740.018.018) and ERC Starting Grant (850818 - MULTIVIsion), as well as an NWO XS grant (OCENW.XS2.033).

## CRediT author statement

**Srividya Ganapathy**: Conceptualization; Methodology; Formal analysis; Investigation; Data Curation; Writing - Original Draft; Visualization. **Xin Meng**: Methodology; Software; Formal analysis; Investigation; Data Curation; Writing - Original Draft; Visualization. **Delizia Mossel**: Formal analysis; Investigation. **Mels Jagt**: Formal analysis; Investigation. **Daan Brinks**: Conceptualization; Formal analysis; Data Curation; Writing - Original Draft; Visualization; Supervision; Project administration; Funding acquisition.

## Data Availability

Data underlying this publication have been deposited in the 4TU.REsearchData repository and are available under DOI 10.4121/21401808

## Supporting Information

**Table S1.**
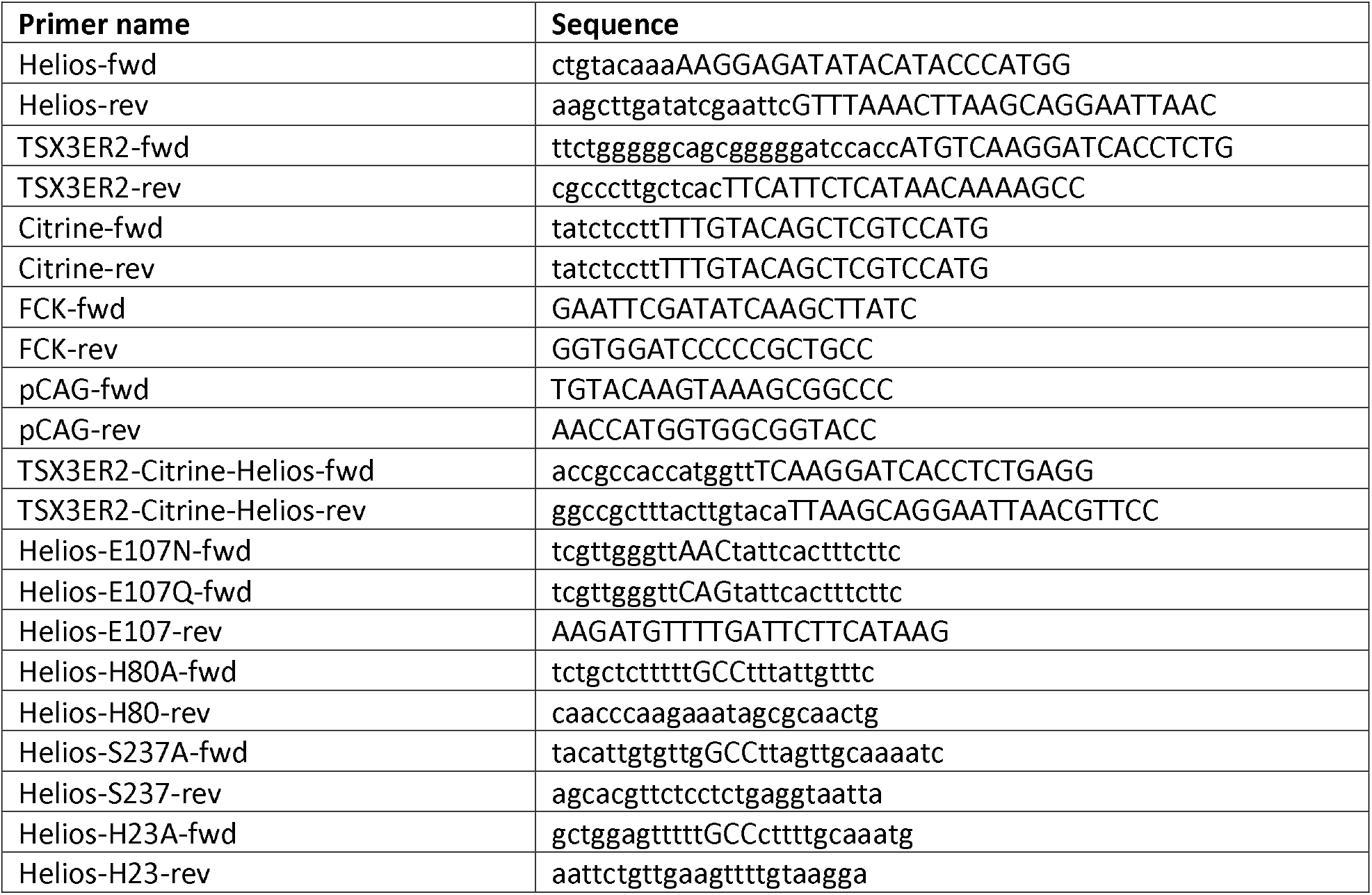
Primer sequences used in this study. Primers used for Gibson assembly cloning of Helios into the expression construct used in this study and for further generating the various site-directed mutants described here.

**Fig. S1.**
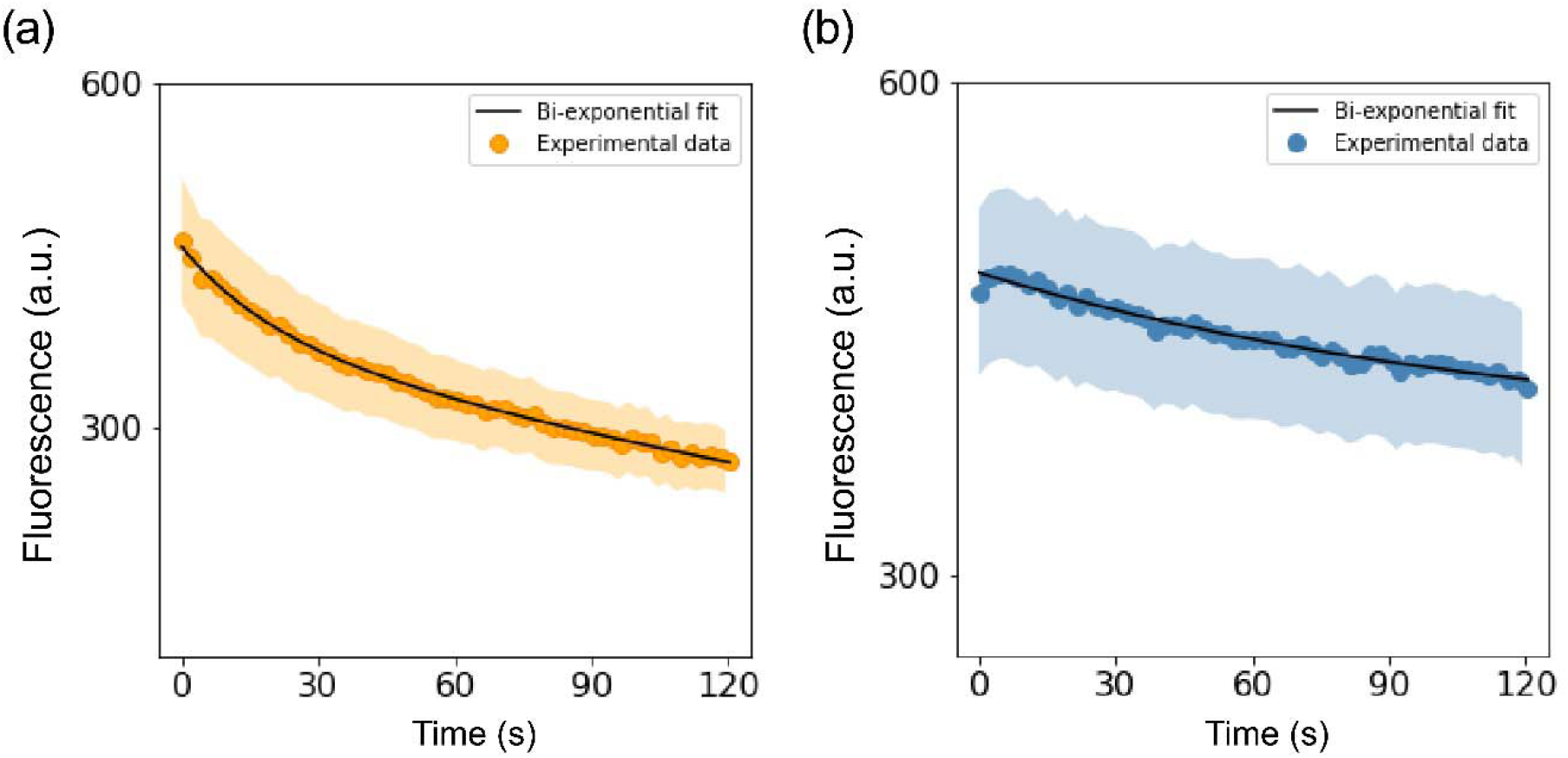
Photobleaching of Helios expressed in *E. coli*. Photobleaching curves of Helios under 561 nm (nm) and 640 nm (b) laser-scanning confocal illumination acquired at a scan speed of 1 fps.

**Fig. S2.**
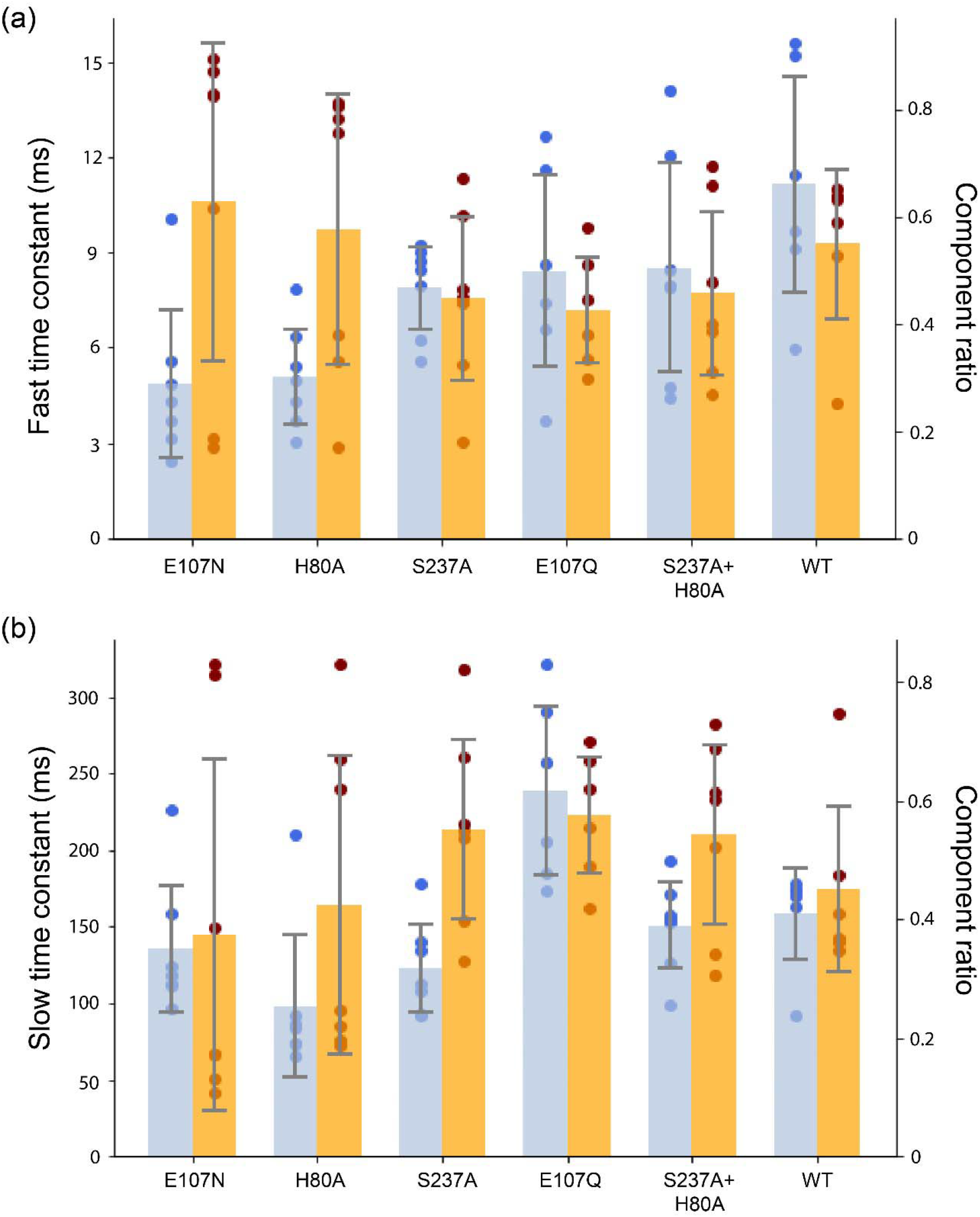
Photobleaching timeconstants and component percentages for 6 Helios mutants. Result bioexponential fits to the photobleaching curves recorded under 87.6 mW/mm2 illumination with 532 nm.

**Fig. S3.**
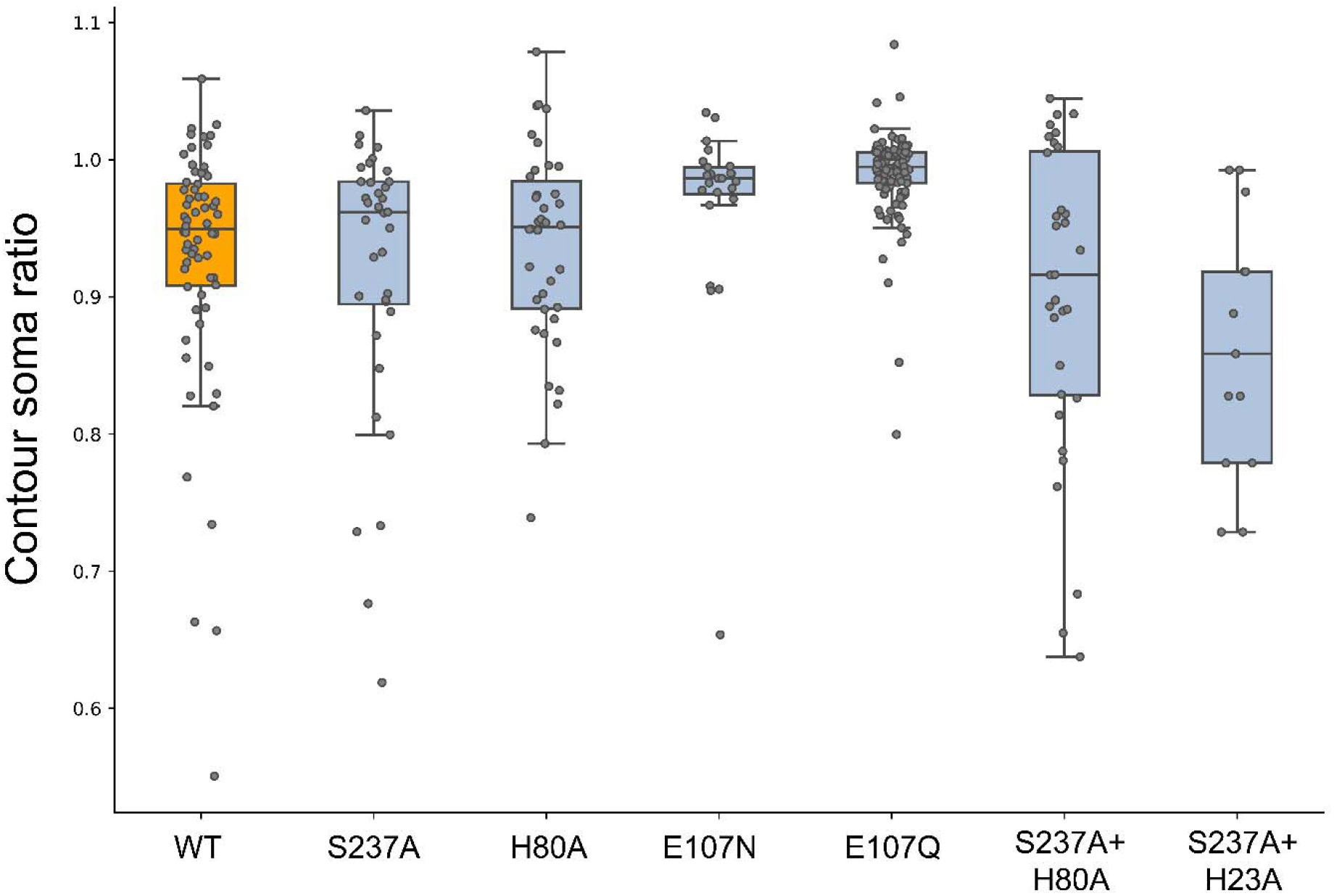
The fluorescence ratios between the cell contour membrane and soma expression from Helios WT and its mutants. The boundaries of the whiskers are based on an interquartile range of 1.5, and each grey point in the plot represents one measurement.

## References

1. Mollinedo-Gajate, I., Song, C. &Knöpfel, T. Genetically Encoded Voltage Indicators. Adv. Exp. Med. Biol. 1293, 209–224 (2021).

2. Gong, Y. The evolving capabilities of rhodopsin-based genetically encoded voltage indicators. Curr Opin Chem Biol 27, 84–89 (2015).

3. Kralj, J. M., Hochbaum, D. R., Douglass, A. D. & Cohen, A. E. Electrical spiking in Escherichia coli probed with a fluorescent voltage-indicating protein. Science (80-). 333, 345–348 (2011).

4. Hochbaum, D. R. et al. All-optical electrophysiology in mammalian neurons using engineered microbial rhodopsins. Nat Methods 11, 825 (2014).

5. McIsaac, R. S. et al. Directed evolution of a far-red fluorescent rhodopsin. Proc Natl Acad Sci U S A 111, 13034–13039 (2014).

6. Saint Clair, E. C. et al. Near-IR Resonance Raman Spectroscopy of Archaerhodopsin 3: Effects of Transmembrane Potential. J Phys Chem B 116, 14592–14601 (2012).

7. Gong, Y., Li, J. Z. & Schnitzer, M. J. Enhanced Archaerhodopsin Fluorescent Protein Voltage Indicators. PLoS One 8, e66959 (2013).

8. Chien, M.-P. et al. Photoactivated voltage imaging in tissue with an archaerhodopsinderived reporter. Sci. Adv. 7, eabe3216 (2021).

9. Piatkevich, K. D. et al. A robotic multidimensional directed evolution approach applied to fluorescent voltage reporters. Nat Chem Biol 14, 352–360 (2018).

10. Tian, H. et al. All-optical electrophysiology with improved genetically encoded voltage indicators reveals interneuron network dynamics in vivo. bioRxiv 2021.11.22.469481 (2021). doi:10.1101/2021.11.22.469481

11. Fan, L. Z. et al. All-Optical Electrophysiology Reveals the Role of Lateral Inhibition in Sensory Processing in Cortical Layer 1. Cell 180, 521–535.e18 (2020).

12. Maclaurin, D., Venkatachalam, V., Lee, H. & Cohen, A. E. Mechanism of voltage-sensitive fluorescence in a microbial rhodopsin. Proc Natl Acad Sci U S A 110, 5939 (2013).

13. Brinks, D., Klein, A. J. & Cohen, A. E. Two-Photon Lifetime Imaging of Voltage Indicating Proteins as a Probe of Absolute Membrane Voltage. Biophys. J. 109, 914–921 (2015).

14. Andreeva, A., Kolev, V. & Lazarova, T. Fluorescence spectroscopy of bacteriorhodopsin at room temperature. https://doi.org/10.1117/12.320967_3573, 359–362 (1998).

15. Kralj, J. M., Douglass, A. D., Hochbaum, D. R., Maclaurin, D. & Cohen, A. E. Optical recording of action potentials in mammalian neurons using a microbial rhodopsin. Nat Methods 9, 90–95 (2012).

16. Flytzanis, N. C. et al. Archaerhodopsin variants with enhanced voltage-sensitive fluorescence in mammalian and Caenorhabditis elegans neurons. Nat Commun 5, 4894 (2014).

17. Pushkarev, A. et al. A distinct abundant group of microbial rhodopsins discovered using functional metagenomics. Nature 558, 595–599 (2018).

18. Hososhima, S. et al. Proton-transporting heliorhodopsins from marine giant viruses. bioRxiv 11, 2022.03.24.485645 (2022).

19. Shihoya, W. et al. Crystal structure of heliorhodopsin. Nature 574, 132–136 (2019). 20.

20. Chazan, A. et al. Diverse heliorhodopsins detected via functional metagenomics in freshwater Actinobacteria, Chloroflexi and Archaea. Environ. Microbiol. 24, 110–121 (2022).

21. Flores-Uribe, J. et al. Heliorhodopsins are absent in diderm (Gram-negative) bacteria: Some thoughts and possible implications for activity. Environ. Microbiol. Rep. 11, 419–424 (2019).

22. Kovalev, K. et al. High-resolution structural insights into the heliorhodopsin family. Proc. Natl. Acad. Sci. 117, 4131 (2020).

23. Singh, M., Inoue, K., Pushkarev, A., Béjà, O. & Kandori, H. Mutation Study of Heliorhodopsin 48C12. Biochemistry 57, 5041–5049 (2018).

24. Meng, X. et al. A compact microscope for voltage imaging. J. Opt. 24, 054004 (2022). 25.

25. Barneschi, L. et al. On the fluorescence enhancement of arch neuronal optogenetic reporters. Nat. Commun. 2022 131 13, 1–9 (2022).

26. Adam, Y. et al. Voltage imaging and optogenetics reveal behaviour-dependent changes in hippocampal dynamics. Nature 569, 413–417 (2019).

27. Hoffmann, M. et al. Color tuning in rhodopsins: the mechanism for the spectral shift between bacteriorhodopsin and sensory rhodopsin II. J Am Chem Soc 128, 10808–10818 (2006).

28. Ernst, O. P. et al. Microbial and animal rhodopsins: structures, functions, and molecular mechanisms. Chem Rev 114, 126–163 (2014).

29. Singh, M., Katayama, K., Béjà, O. & Kandori, H. Anion binding to mutants of the Schiff base counterion in heliorhodopsin 48C12. Phys. Chem. Chem. Phys. 21, 23663–23671 (2019).

30. Bada Juarez, J. F. et al. Structures of the archaerhodopsin-3 transporter reveal that disordering of internal water networks underpins receptor sensitization. Nat. Commun. 2021 121 12, 1–10 (2021).

31. Tomida, S., Kitagawa, S., Kandori, H. & Furutani, Y. Inverse Hydrogen-Bonding Change between the Protonated Retinal Schiff Base and Water Molecules upon Photoisomerization in Heliorhodopsin 48C12. J. Phys. Chem. B 125, 8331–8341 (2021).

32. Silapetere, A. et al. QuasAr Odyssey: The Origin of Fluorescence and its Voltage Sensitivity in Microbial Rhodopsins. Rev. Process Nat. Commun. 13, 1–20 (2022).

33. Wijesiri, K. &Gascón, J. A. Microsolvation Effects in the Spectral Tuning of Heliorhodopsin. J. Phys. Chem. B 126, 5803–5809 (2022).

34. Besaw, J. E. et al. Low pH structure of heliorhodopsin reveals chloride binding site and intramolecular signaling pathway. Sci. Reports 2022 121 12, 1–16 (2022).

